# The variable domain from the mitochondrial fission mechanoenzyme Drp1 promotes liquid-liquid phase separation

**DOI:** 10.1101/2023.05.29.542732

**Authors:** Ammon E. Posey, Mehran Bagheri, Kyle A. Ross, Elizabeth N. Lanum, Misha A. Khan, Christine M. Jennings, Megan C. Harwig, Nolan W. Kennedy, Vincent J. Hilser, James L. Harden, R. Blake Hill

## Abstract

Dynamins are an essential superfamily of mechanoenzymes that remodel membranes and often contain a “variable domain” (VD) important for regulation. For the mitochondrial fission dynamin, Drp1, a regulatory role for the VD is demonstrated by mutations that can elongate, or fragment, mitochondria. How the VD encodes inhibitory and stimulatory activity is unclear. Here, isolated VD is shown to be intrinsically disordered (ID) yet undergoes a cooperative transition in the stabilizing osmolyte TMAO. However, the TMAO stabilized state is not folded and surprisingly appears as a condensed state. Other co-solutes including known molecular crowder Ficoll PM 70, also induce a condensed state. Fluorescence recovery after photobleaching experiments reveal this state to be liquid-like indicating the VD undergoes a liquid-liquid phase separation under crowding conditions. These crowding conditions also enhance binding to cardiolipin, a mitochondrial lipid, raising the possibility that phase separation may enable rapid tuning of Drp1 assembly necessary for fission.

## Introduction

Members of the dynamin superfamily are multi-domain GTPases that utilize GTP hydrolysis to either divide or fuse cellular membranes (1, 2). These proteins share a specific domain architecture that includes, at a minimum, a G domain that binds GTP with low affinity and hydrolyzes it to GDP, and a stalk domain that mediates oligomerization in some family members and self-assembly in others (3, 4) (**Fig. 1A**). Self-assembly enhances GTP hydrolysis as demonstrated by the prototypical dynamin, dynamin-1, where assembly can stimulate hydrolysis 100-fold (5–7). Dynamin family members may also contain an additional domain involved in membrane targeting. These domains are known as variable domains (VDs) because they vary between family members. For dynamin-1, the VD is a Pleckstrin Homology (PH) domain that binds phosphoinositides aiding plasma membrane localization (8). For the mitochondrial dynamin, Drp1, the VD is not a PH domain, but binds cardiolipin aiding mitochondrial localization, proper assembly, and activity (9–15).

**Figure 1.**
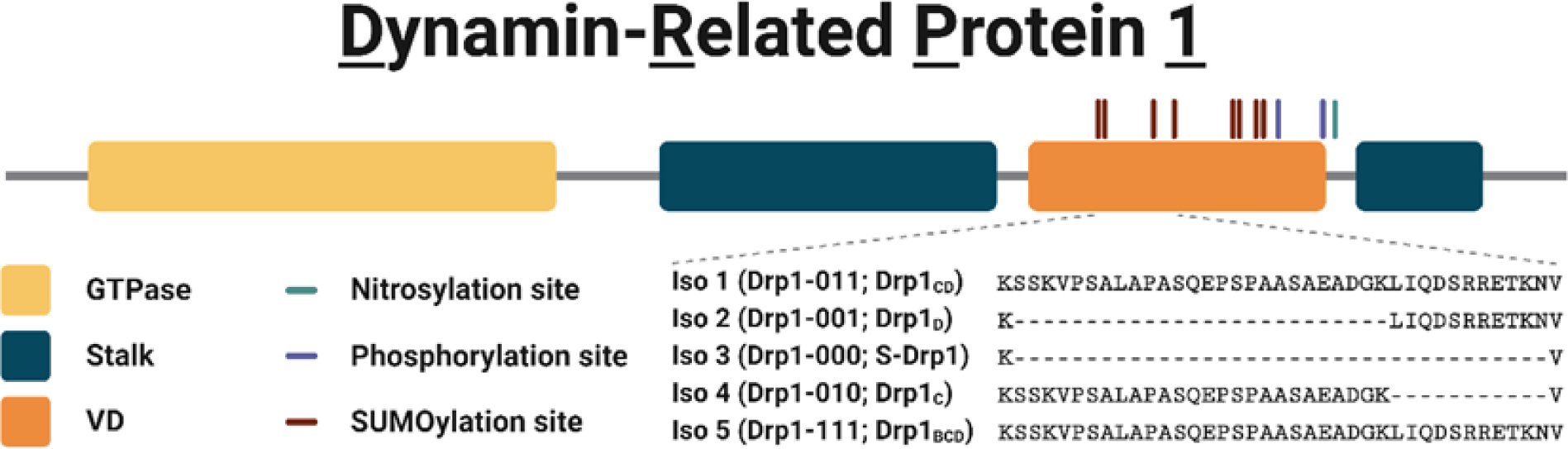
Schematic showing the domain architecture of Drp1 and a subset of the isoforms and post-translational modifications involving the variable domain (VD). The protein construct used in this study is variable domain from isoform 1 (VD1), entailing residues 501-637. Created with BioRender.com.

The VDs of dynamins appear exquisitely linked to enzyme function, which is perhaps most apparent for Drp1. The VD of Drp1 harbors two phosphorylation sites (16–20) and eight SUMOylation sites (21–24). These post-translational modifications are able to activate or inhibit mitochondrial fission consistent with the VD governing Drp1 activity (25). Supporting this model, Drp1 is differentially expressed in human tissues as different isoforms (26–29), five of which differ by alternative splicing in the VD (30). Deletions and substitutions in the VD paradoxically inhibit or enhance mitochondrial fission (31). In these experiments, expression of Drp1 lacking the VD (Drp1ΔVD) in HeLa cells fragment mitochondria, presumably from increased mitochondrial fission. How Drp1 without the VD can increase fission is unclear because the VD mediates mitochondrial targeting through binding to the mitochondrial-specific lipid cardiolipin (10, 12, 14, 32). Further, VD-mediated membrane association enhances GTPase activity (11, 12, 33–35). The model that emerges from these data suggests that the VD inhibits assembly of Drp1 in solution, and that inhibition is relieved upon membrane binding (**Fig. 1B**).

In contrast to this model, several data support a more complex role for VD in modulating Drp1 activity. Drp1 lacking the VD is incapable of tubulating spherical lipid vesicles (11, 34, 35) yet is able to bind membranes (11) suggesting that the VD is not the sole determinant of membrane-stimulated hydrolysis. Indeed, the stalk domain of Drp1 alone also binds membranes (36); further, bioinformatic considerations implicate a conserved region of the GTPase domain in membrane binding (37). These data suggest that VD contributes to, but is not essential for, membrane binding. However, transient VD-membrane interactions also induce cardiolipin clustering that can form condensed membrane regions important for fission (11). Furthermore, the alterations to the VD dramatically alter Drp1 localization. Drp1 inside cells is found by fluorescence microscopy in two populations: a diffuse haze throughout the cytoplasm and a subset in bright dots (punctate foci, or puncta), that often, but not always, co-localize with mitochondria. Drp1 foci are thought to be pre-scission complexes comprised of assembled Drp1 poised for membrane scission. However, previous FRAP work shows rapid recovery upon photobleaching suggesting a less well-ordered state. Mutational analysis of the VD identified both gain- and loss-of-function variants with dramatic changes in punctate structures and cellular distributions in a manner that suggests something more than Drp1 assembly is impaired when the VD is mutated (31). Taken together these results suggest that the Drp1 VD plays multiple roles in regulating activity, but the basis for this is still unclear.

The VD is intrinsically disordered by prediction and circular dichroism (12), consistent with frequent occurrence of disordered regions in regulatory hubs that are sites of alternative splicing and post-translational modifications (38). To gain insight into intrinsic VD properties, we expressed and purified the VD from Drp1 isoform 1 (VD1) and determined by biophysical methods that VD1 is indeed intrinsically disordered throughout the polypeptide chain. Surprisingly, under crowding conditions similar to that on the membrane surface, VD1 condenses into a separate liquid phase, which appears to promote membrane interactions. These findings identify a novel property of the VD that may help explain the aberrant morphologies discovered earlier upon mutation to the VD and suggest that native Drp1 foci in cells may arise, in part, from phase separation that may be modulated by the known alterations to VD.

## Results

### The Drp1 variable domain is intrinsically disordered

A variable domain construct composed of residues 501-637 of Drp1 isoform 1 (VD1) was expressed and purified for structural analyses. The protein was well-expressed into the soluble fraction of E. coli with a yield of ∼15 mg homogeneous protein/L rich media. When analyzed by size exclusion chromatography (SEC), the VD1 eluted much earlier than would be expected for a globular protein of its molecular weight (**Fig. 2A**). Using a standard curve composed of the S75 elution volumes of eight globular proteins of known molecular weight, we calculated an expected elution volume of 89 mL for a globular monomer with a molecular weight equivalent to that of the VD1 construct (15.4 kDa). The observed elution volume for the VD1 was 69 mL, which based on the standard curve equates to an apparent molecular weight that is approximately 3.5 times larger than that expected for a monomer (assuming globular shape). The results may be interpreted to mean that VD1 forms globular oligomers, a result that was previously reported for a VD construct (39). However, an alternative interpretation is the lower elution volume is consistent with dimeric or monomeric extended chains with a large hydrodynamic radius.

**Figure 2.**
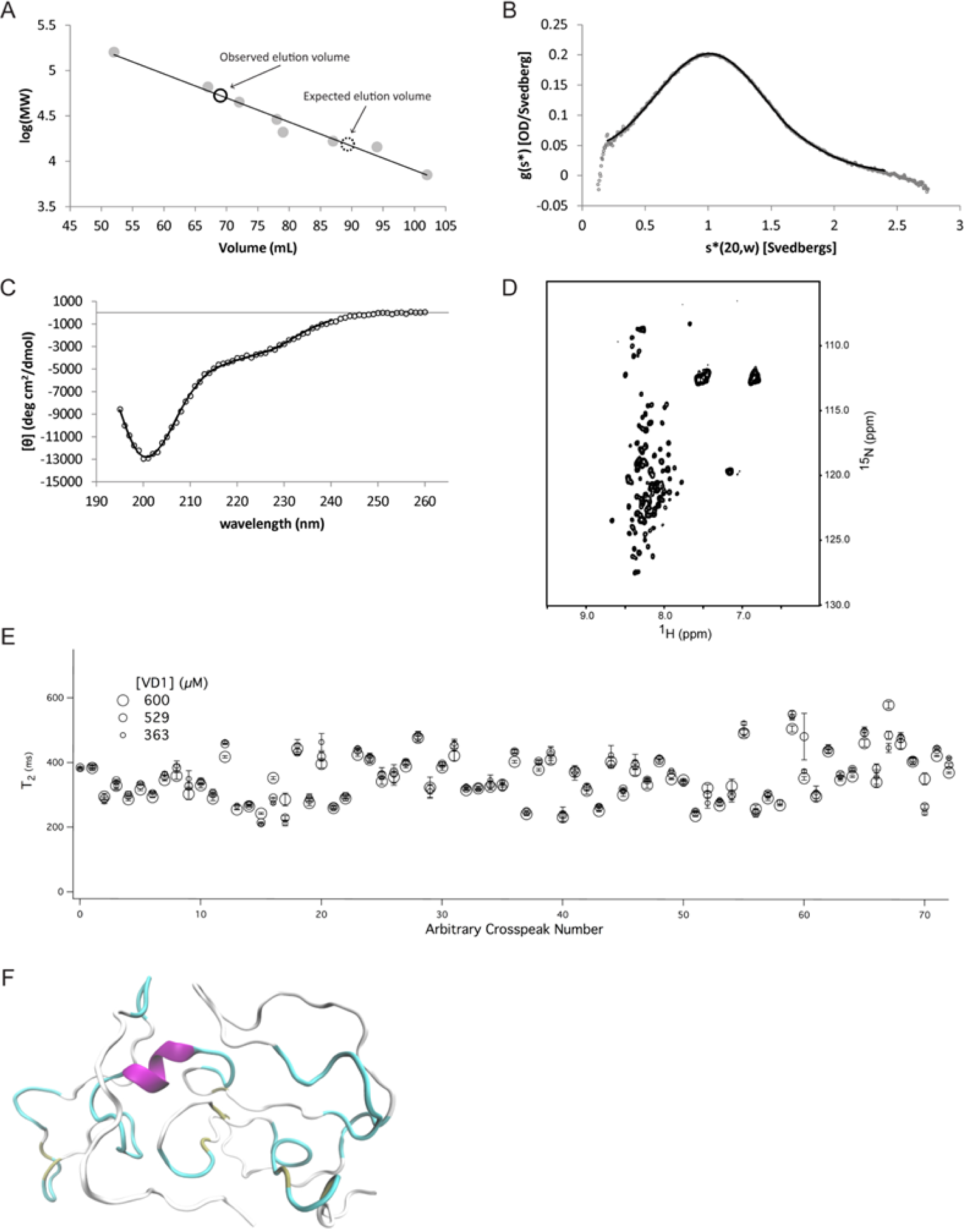
Structural properties of the isolated variable domain from Drp1. **A**, VD1 eluted from an S75 gel filtration column at ∼69 mL (open solid black circle), much earlier than the 89 mL elution volume for a globular protein of equivalent molecular weight (open dashed black circle) based on globular protein standards (filled gray circles). **B**, Fit of sedimentation velocity data of VD1 to the Lamm equation using DCDT+ gives a sedimentation coefficient of 1.069 S_20,w_. **C**, Circular dichroism spectrum of the VD1 is consistent with intrinsic disorder, with a characteristic minimum near 200 nm. **D**, a ^1^H-^15^N HSQC spectrum of 300 µM VD1 collected at 18.8 T in Buffer A, 25 °C. **E**, Spin-spin relaxation time constants (T_2_) were collected on VD1 samples at the indicated concentrations at 18.8T **F**, Representative conformation of VD1 taken from the last 50ns of a 300-ns molecular dynamics simulation of the variable domain in water showing backbone structure with alpha-helix (purple), coil (white), and turn (cyan). All measurements were conducted in Buffer A at 25 °C.

To test this hypothesis, we turned to sedimentation velocity (**Fig. 2B**), which indicated that the VD1 is indeed predominantly monomeric with a sedimentation coefficient of 1.069 S_20,w_ and an estimated molecular weight of 13,970 Da (vs. 15,363 Da calculated from sequence). From the sedimentation velocity data, the maximum sedimentation coefficient was estimated to be S_max_ = 2.231, and represents the sedimentation coefficient of the smallest non-hydrated smooth sphere that could contain the mass of a given protein. By taking the ratio of S_max_ and the measured S_20,w_, a S_max_/S_20,w_ ratio of 2.09 is obtained and indicates that VD1 populates moderate to highly elongated conformations as globular proteins typically have a ratio near 1.2 or 1.3, moderately elongated proteins have a ratio around 1.5 to 1.9, and highly elongated proteins have a ratio around 2.0 to 3.0 (40). Comparison of the measured VD1 hydrodynamic dimensions to those published for other globular, pre-molten globule, random coil and denatured proteins (39) indicated that the hydrodynamic dimensions of the VD1 lie between those of a natively unfolded pre-molten globule and a natively unfolded random coil (**Fig. S1**).

To differentiate between these conformations, the secondary structure of VD1 was evaluated by circular dichroism (CD) (**Fig. 2C**). A slight minimum at 222 nm and a more profound minimum near 200 nm that was temperature independent indicate a largely unfolded conformation with the possibility of some α-helix, similar to that observed for VD3, the shorter variable domain from Drp1 isoform 3 (12). A comparison of CD data to other proteins (41) is consistent with VD1 adopting a pre-molten globule-like conformation (**Fig. S1**). A ^1^H-^15^N HSQC spectrum of uniformly ^15^N-labeled VD1 showed poor chemical shift dispersion, especially in the ^1^H dimension with resonances centered at random coil chemical shift values (**Fig. 2D**). We also measured NMR spin-spin relaxation rates at three concentrations (363, 529, and 600 µM) and found little evidence of concentration-dependent decreases in T_2_ that would be indicative of self-association (**Fig. 2E**). Thus, VD1 exists as a monomer under these conditions. To assess whether VD1 might adopt fleeting amounts of secondary structure consistent with a pre-molten globule-like conformation, we conducted MD simulations of VD1 in explicit water. Starting with an elongated chain, the radius of gyration (R_g_) reached a plateau after 250ns with an average R_g_ = 17.2 ± 0.3 Å. The steady-state VD1 conformations display somewhat elongated globular configurations consistent with our hydrodynamic measurements. We found little evidence of stable secondary structural elements, although fleeting amounts of α-helix (∼7%) persisted especially for residues 507-517 and 580-588 (**Fig. 2F**) consistent with our CD data. Collectively, these data definitively indicate that the VD1 is intrinsically disordered similar to the isolated variable domain from isoform 3 (42).

### The VD1 undergoes an apparently two-state transition in the protecting osmolyte TMAO

Since VD1 is intrinsically disordered and is proposed to play an autoinhibitory role, we sought to determine if VD1 adopts a latently folded/ordered conformation consistent with a disordered-to-ordered transition that might help relieve autoinhibition as in other systems (43–45). We used the protecting osmolyte trimethylamine-N-oxide (TMAO) that is known to induce folding of proteins with destabilized or latent folds (46–49). TMAO is a naturally occurring osmolyte known to favor native-like folded states of proteins by elevating the free energy of the unfolded state (47). To determine the effect of TMAO on VD1, we followed the intrinsic fluorescence of the single tryptophan at position 89 of the isolated VD1 construct (589 in Drp1 iso 1) in absence and presence of 4.35 M TMAO (**Fig. 3A**). The wavelength of maximum tryptophan fluorescence intensity (λ_*max*_) shifted significantly to the blue by 12 nm, a strong indicator of tryptophan burial. (**Fig. 3A-B**, **filled circles**). The fluorescence intensity measured at a single wavelength (338 nm) as a function of increasing TMAO concentration showed a cooperative transition with a midpoint coincident with λ_*max*_ (**Fig. 3B**). This transition was not attributable to a change in solvent polarity from TMAO as titrations with free indole showed a negligible transition with a red shift of 4 nm in λ_*max*_ (data not shown). The VD1 data are well fit to two-state folding model (**Fig. 3B**, **solid line**) giving an m-value of 2.9 ± 0.2 kcal•mol^-1^M^-1^, which is similar to TMAO-induced folding of other intrinsically disordered proteins of this size (50–53).

**Figure 3.**
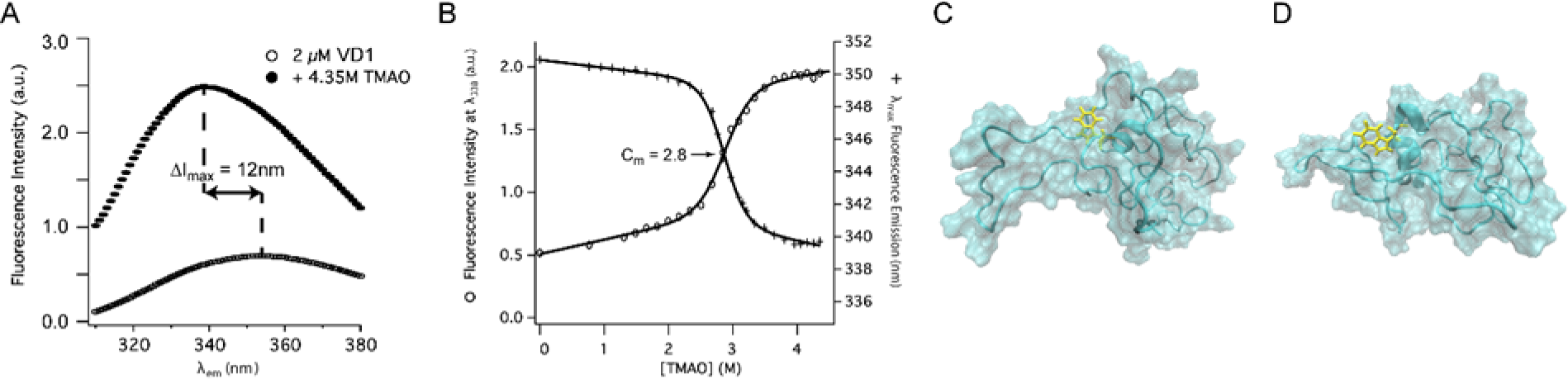
Steady-state tryptophan fluorescence of VD1 in the protecting osmolyte TMAO. ***A***, Intrinsic tryptophan fluorescence emission spectrum of 2 µM variable domain in the absence (lower trace, open circles) and presence (upper trace, filled circles) of TMAO. Note that the addition of the protective osmolyte TMAO increased the fluorescence intensity and blue-shifted λ_max_ by 12 nm. ***B***, Fluorescence intensity (open circles) and λ_max_ (plus signs) of 2 µM variable domain at 338 nm as a function of TMAO fit to a two-state model (solid line). Note the coincidence in the C_m_ for both measurements. ***C-D,*** Representative conformations of the last 50 ns from a 300ns molecular dynamics simulation of VD1 in water (left) and TMAO (right).

Since the m-value correlates with the amount of protein surface buried from solvent upon folding (54), these data suggest TMAO induces VD1 folding. To assess this, we attempted to measure CD, but light scattering from TMAO prevented reliable ellipticity measurements. Insight into the TMAO-induced state, was gleaned by conducting a 300 ns MD simulation in the presence of 3.3 M TMAO. We found little indication of tryptophan burial or folding, although a small increase in transient α-helix was observed (**Fig. 3C-D**). These data suggest the change in tryptophan fluorescence observed upon TMAO addition might not arise from folding.

### The TMAO-induced transition involves self-association

In preparing the above samples, we noted that the VD1 stock solution in TMAO was noticeably turbid, but the turbidity vanished upon 15x dilution to the final protein concentration. This reversible turbidity at higher protein concentrations in the presence of TMAO could be indicative of molecular self-association. To assess this, static right-angle light scattering (RALS) of the same samples used for fluorescence measurements was collected. A cooperative transition was observed with an identical C_m_ of 2.8 M (**Fig. 4A**) that was essentially superimposable on the fluorescence data in both the forward and reverse (not shown) titrations. This suggests that fluorescence and light scattering are reporting on the same process. As a control, we also measured RALS as a function of TMAO for lysozyme, which is similar to VD1 in molecular weight, but is natively folded. Lysozyme showed no sign of self-association (as measured by light scattering) even at high concentrations of TMAO (**Fig. S2**).

**Figure 4.**
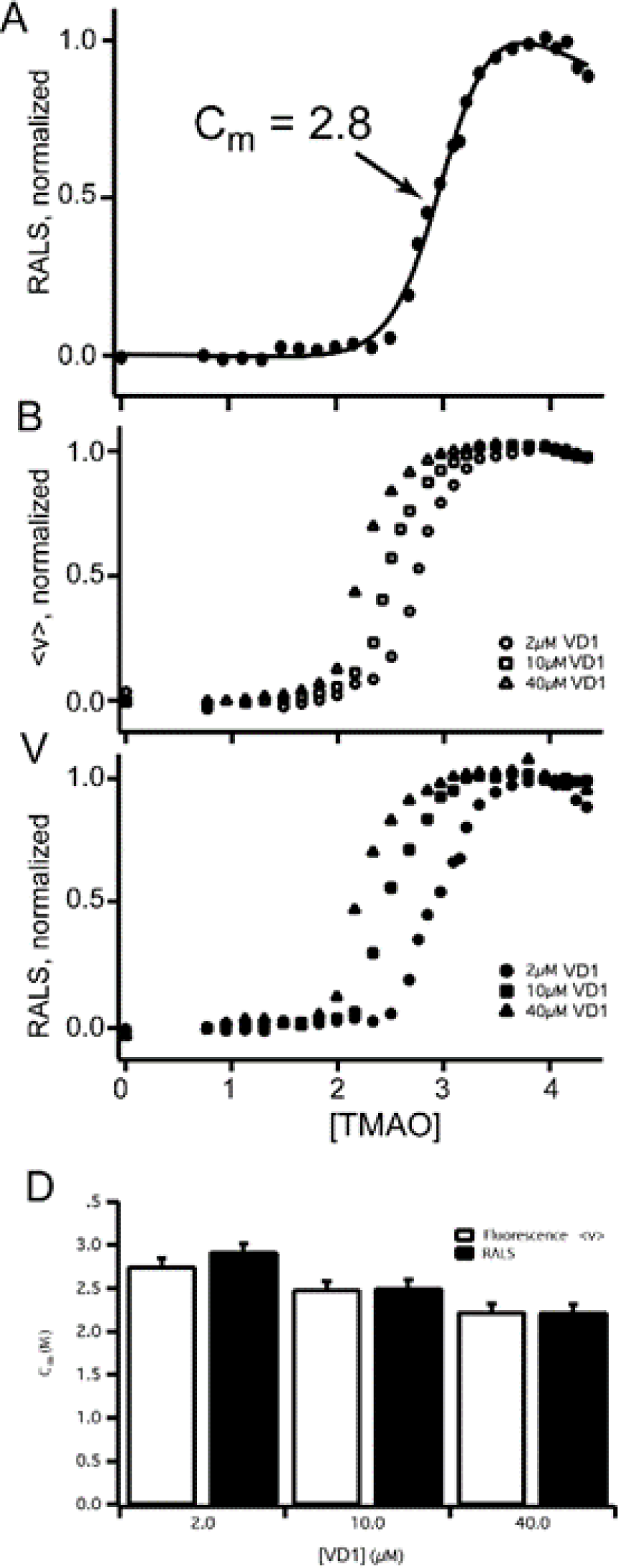
Concentration dependence of VD1 in the protecting osmolyte TMAO by light scattering and fluorescence. ***A***, 90° light scattering data (filled circles) of 2µM variable domain at 450 nm as a function of TMAO fit to a two-state model (solid line) with a C_m_ = 2.8. ***B-D***, Concentration dependence to the TMAO-induced tryptophan fluorescence transition of the variable domain at 2 μM (circles), 10 μM (squares) and 40 μM (triangles). ***D***, The same samples as in (***B***) followed by (***C***) light scattering (filled shapes) give identical TMAO titration midpoints, C_m_.

If TMAO-induced light scattering of VD1 requires self-association, then it should be concentration-dependent. To test this, we repeated the TMAO titrations at 2, 10, and 40 µM VD1. At each protein concentration, both fluorescence and light scattering gave identical transitions and midpoints (**Fig. 4B-D**) with a clear dependence on protein concentration. By contrast, VD1 shows no concentration-dependent change in tryptophan λ*_max_* in the absence of TMAO (**Fig. S3**) indicating that VD1 self-association requires TMAO. In these experiments the changes in turbidity were entirely reversible upon dilution. We conclude that VD1 undergoes a reversible and cooperative TMAO-induced self-association that creates assemblies sufficiently large to scatter light. This process does not appear to involve cooperative folding of VD1.

### VD1 self-association in the presence of TMAO is consistent with a phase transition

A notable feature of the VD1 in TMAO is the reversible turbidity that suggested a directed assembly as opposed to an amorphous aggregation. To address this, we used transmission electron microscopy (TEM). The same samples used for fluorescence and RALS measurements were applied to grids and imaged by TEM. The TMAO-induced state observed by TEM was neither an amorphous aggregate nor fibril-like ordered assembly. Rather, a darkly stained spherical or droplet-like assemblies (**Fig. 5A-B**), 100-200 nm in diameter, was consistently observed with negatively stained globular textures within. This morphology was not observed in the TMAO-only or protein-only controls. Samples prepared minutes to hours before being applied to the grid showed such assemblies, which appeared to be increasingly abundant in samples that were incubated in TMAO for longer periods (many hours to days). Interfacial tension between two immiscible liquid phases drives the minimization of the interfacial surface area between the phases. Because a sphere achieves the desired minimal surface area, spherical droplets are formed (55). Thus, VD1 assemblies in TMAO with spherical morphology are suggestive of a liquid-liquid phase transition.

**Figure 5.**
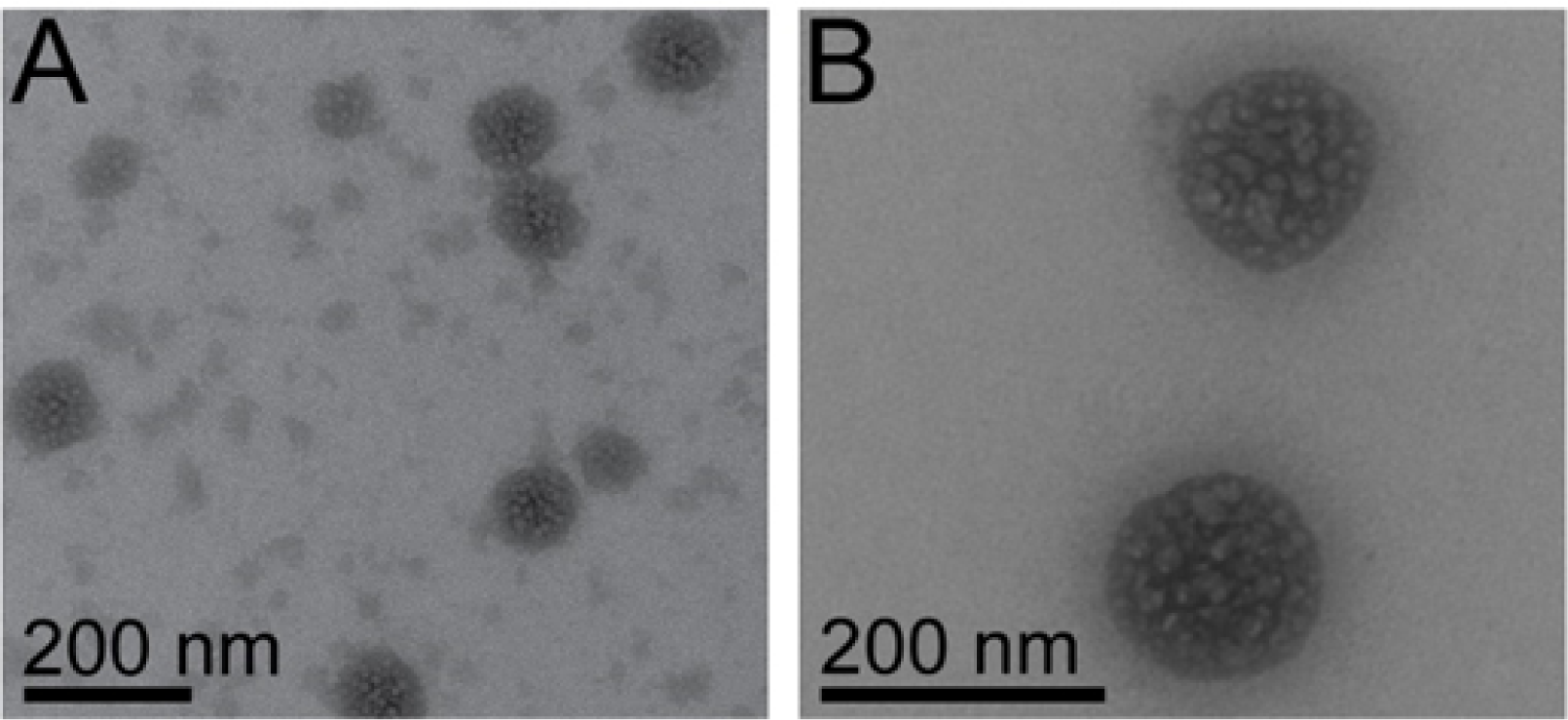
TMAO induces phase separation of the Drp1 VD1. ***A-B***, Transmission electron microscopic images of 2 μM variable domain in 3 M TMAO. The two samples were independently prepared under the same conditions and are representative of several observations. The darkly-stained spherical phase separates are roughly 50 – 150 nm in diameter and appear to be formed by the coalescence of smaller globules (negatively-stained texture within the larger spheres). Scale bar is 200 nm in both images.

### VD1 separates into a distinct phase in the presence of other solutes

Expanding on the reversible, TMAO-induced phase separation, we investigated whether other cosolutes might induce a phase-separated state with VD1. For this, we used ammonium sulfate (NH4)_2_SO_4_, polyethylene glycol PEG-8000 and Ficoll PM 70, which are known to induce condensed states in a variety of systems (56–61). Light scattering of increasing concentrations of VD1 were monitored for increased turbidity (**Fig. 6A**). The addition of ammonium sulfate to VD1 triggered an instantaneous response in which the mixture became very turbid. VD1 titrations with 20% PEG-8000 or 20% Ficoll PM 70 also produced increased light scattering in a concentration-dependent manner (**Fig. 6A**), but to a lesser extent. In each case, turbidity was reversible. By differential interference contrast (DIC) microscopy, all conditions with VD1 caused two types of condensed states: a spherical droplet phase that was free-floating and a much less spherical phase that was associated with the bottom of the well (representative images, **Fig. 6B-D**). Pre-treating of wells with detergents or surfactants did not alter the formation of this “base layer”, which appeared in a time- and concentration-dependent manner as did the free-floating droplets (**Fig. S6)**. In other systems, weak ionic interactions help support phase separation (62–64), which we assessed by fluorescence microscopy with an AlexaFluor-labeled VD1 and increasing potassium chloride in the presence of 20% PEG-8000. Increasing salt decreased the propensity to phase separate which was quantified by determining the coefficient of variation of the fluorescence intensity signal across the entire microscope field (**Fig. 6E**). For solutions that do not form droplets, the coefficient of variation is close to zero, whereas droplet formation creates regions of high intensity that increases the coefficient of variation. These data are a subset of several experiments in which different concentrations of protein, solute, and salt modulate the ability of VD1 to phase separate.

**Figure 6.**
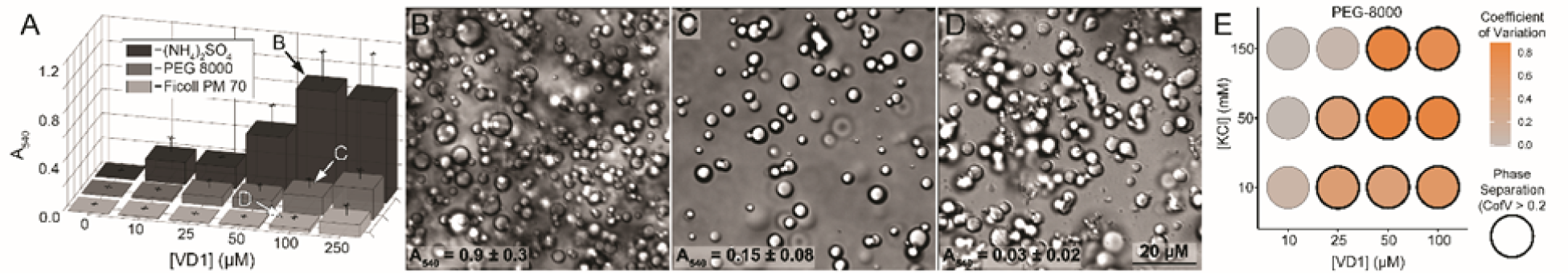
Phase separation of Drp1 VD1 in molecular crowders. ***A***, Light scattering data of 100 µM VD1 at 540 nm as a function of 1.68 M (NH_4_)_2_SO_4_ (black), 20% PEG 8000 (dark grey), and 20% Ficoll PM 70 (light grey). ***B-D***, DIC microscopy images of µM VD1 in 1.68 M (NH_4_)_2_SO_4_ (***B***), 20% PEG 8000 (***C***), and 20% Ficoll PM 70 (***D***). Scale bar = 20 µm. ***E***, Phase diagram of VD1 in PEG 8000 under varying conditions. Black circle outlines indicate LLPS. Orange-grey color gradient within circles indicates the coefficient of variation value calculated from representative images following titration of potassium chloride. Experiments conducted in 20 mM HEPES pH 7.4, 1 mM beta-mercaptoethanol at 25°C.

### Droplets formed by VD1 are highly mobile by FRAP

To determine the nature of the phase-separated state, we used fluorescence recovery after photobleaching (FRAP) that is well-established to discern the overall fluidity of molecules within condensates (65–68). Under conditions that favor the phase-separated state with an AlexaFluor-labeled VD1, droplets were photobleached and recovery intensities monitored (**Fig. 7A-B**). For droplets in PEG-8000 and Ficoll PM 70, fluorescence intensities returned to over 90% of their baseline intensities on average (**Fig. 7C**), indicating high molecular mobility consistent with liquid-like phase separation (LLPS). Droplets formed in ammonium sulfate displayed significantly lower fluorescence recoveries on the same time scale, suggesting that VD1 assemblies within these droplets may adopt a mix of more well-ordered structures. Regardless, VD1 appears capable of forming both liquid-like and gel-like assembly states depending on the environment.

**Figure 7.**
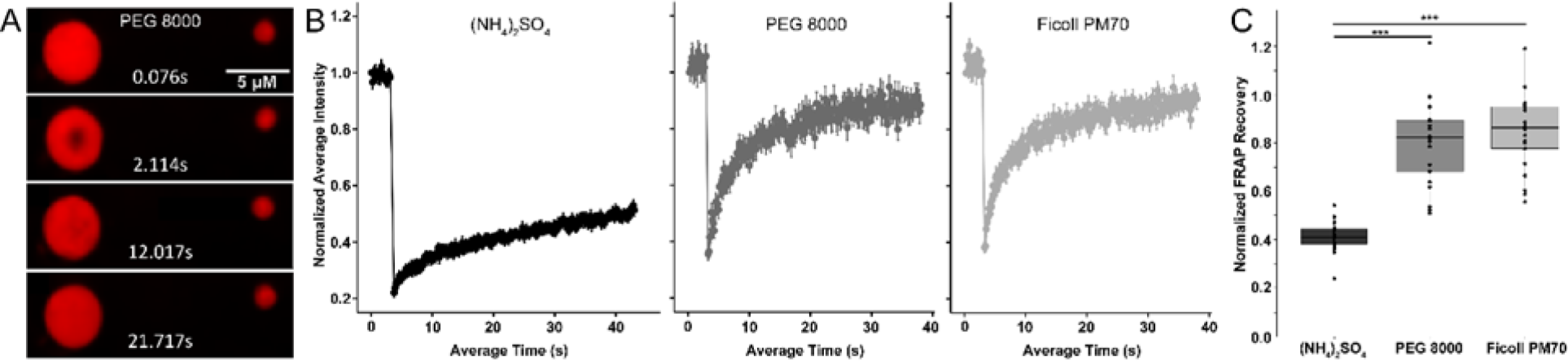
Drp1 VD1 droplets are liquid-like by FRAP. ***A***, Example fluorescence recovery after photobleaching (FRAP) images of 100 µM VD1 in PEG 8000 at various time points. Peripheral droplet included as a reference point for non-specific photobleaching. Scale bar = 5 µm. ***B***, Normalized FRAP intensity curves for 100 µM VD1 in 1.68 M (NH_4_)_2_SO_4_ (black), 20% PEG 8000 (dark grey), and 20% Ficoll PM 70 (light grey). Photobleaching laser power for droplets in (NH_4_)_2_SO_4_, PEG 8000, and Ficoll PM 70 were 15%, 30%, and 100%, respectively. ***C***, FRAP recovery values for VD1 droplets from ***B*** in 1.68M (NH_4_)_2_SO_4_ (black), 20% PEG 8000 (dark grey), and 20% Ficoll PM 70 (light grey). FRAP recovery was calculated as a ratio of the final droplet fluorescence intensity to the initial droplet fluorescence intensity, correcting for the fluorescence intensity after bleach stimulation as follows: 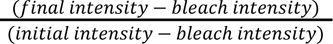. Anova, *p* < 3. 48 × 10^-11^ (one-way ANOVA with Tukey’s post hoc multiple comparisons test, *** *p* < 0. 0001, *n* = 20, 21, or 21 droplets for (NH_4_)_2_SO_4_, PEG 8000, or Ficoll PM 70, respectively). Experiments conducted in 20 mM HEPES pH 7.4, 1 mM beta-mercaptoethanol at 25°C.

### VD1 lipid binding and LLPS is enhanced in the presence of TMAO

Arg/Lys are known to mediate binding to anionic lipids, and this fact with the reduced solvation of VD1 by TMAO (and also ammonium sulfate, **Fig. S5**) caused us to speculate whether TMAO-induced crowding might be akin to the environment that VD1 experiences upon Drp1 assembly at the anionic membrane surface of mitochondria. VD1 preferentially binds cardiolipin (42), a lipid unique to mitochondria, and we reasoned that TMAO-induced desolvation might favor VD1 binding to cardiolipin. To test this idea, VD1 binding to large unilamellar vesicles was measured in a sedimentation assay. The fraction of VD1 that partitioned onto 25% cardiolipin vesicles was ∼0.2 in the absence of TMAO consistent with earlier work (10,12,14,30). By contrast, under conditions that favor a condensed state with TMAO, the fraction of VD1 bound increased four-fold to ∼0.8. Accordingly, the partition coefficient increased from *K_x_*= 3. 2×10 in the absence of TMAO to *K_x_*= 6. 7×10 in the presence of 1.8 M TMAO (**Fig. 8A**). This effect was greater for cardiolipin than other phospholipids tested such as phosphatidylserine and phosphatidyl choline. We interpret the enhanced lipid binding and intact specificity for cardiolipin in the presence of TMAO to verify that the TMAO-condensed state is competent for cardiolipin binding.

**Figure 8.**
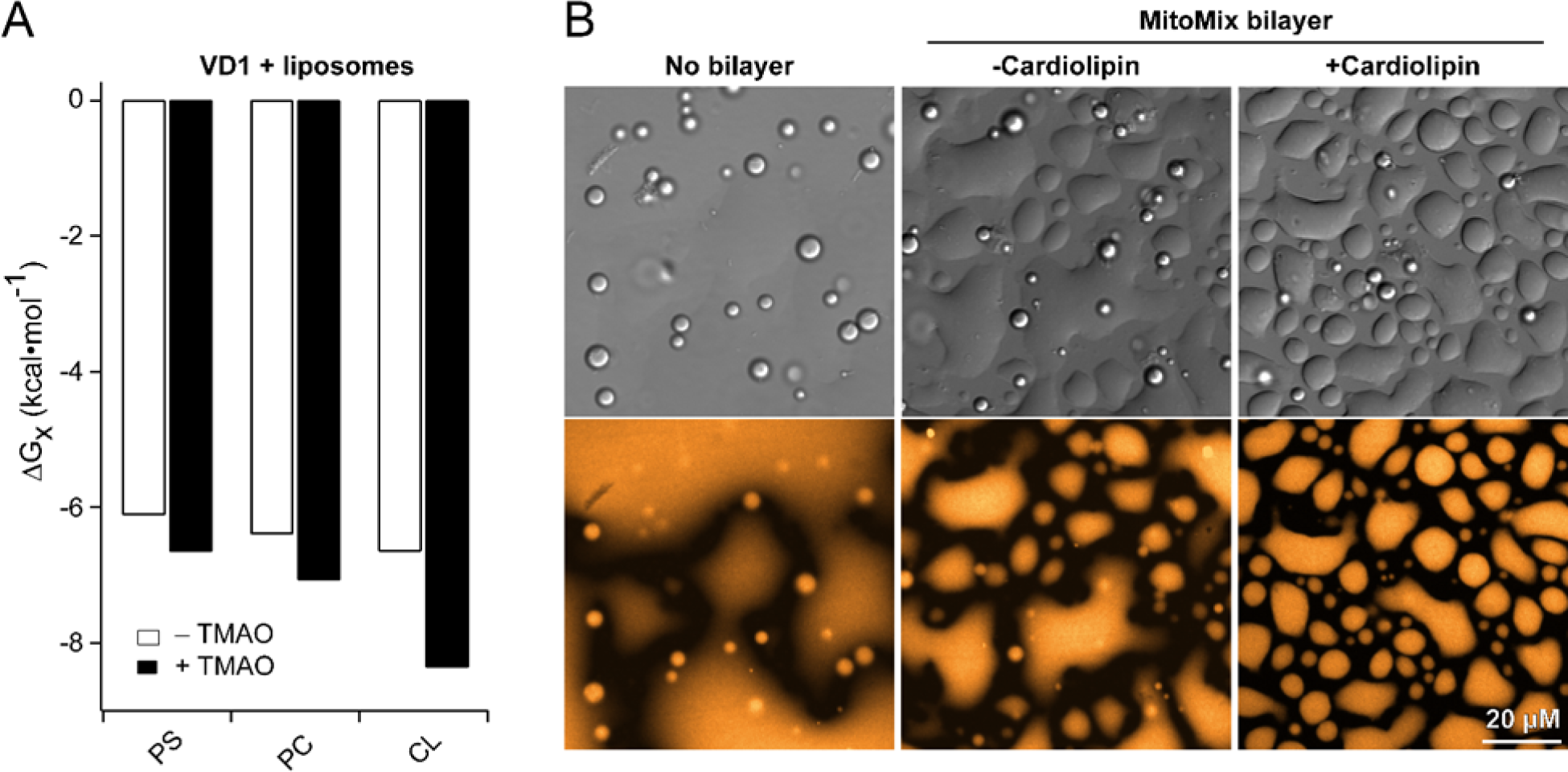
Cardiolipin enhances VD1-membrane interactions and phase separation. ***A***, Membrane binding measurements of the variable domain in the presence of 1.8 M TMAO. Free energies of partitioning were calculated from sedimentation data of the variable domain with 100 nm large unilamellar vesicles (75% DOPC/25% indicated lipid – either phosphatidylserine PS, phosphatidyl choline PC, or cardiolipin CL) at a protein:lipid ratio of 1:200. ***B***, DIC and fluorescence microscopy images of 50 µM VD1 in 20% PEG 8000 on uncoated (left panel), MitoMix bilayer without cardiolipin (middle panel), or MitoMix bilayer with cardiolipin (right panel) coated wells. All measurements at 25 °C in 10 mM HEPES pH 7.4, 50 mM KCl.

These data suggested that a VD1/cardiolipin interaction might modulate LLPS. To test this, supported lipid bilayers with or without cardiolipin were incubated with VD1 at 20% PEG-8000, conditions that promote LLPS (**Fig. 8B, left panel**). In the absence of supported lipid bilayers, DIC microscopy showed spherical droplets formed in 20% PEG-8000 as before, whereas fluorescence microscopy indicated some background fluorescence associated with bottom of the well where VD1 appears to be wetting the surface as a low-surface tension liquid. Under identical conditions, the addition of supported lipid bilayers also caused spherical, free-floating droplets similar to the absence of bilayers, and appeared to enhance the “base layer” phase that was much less spherical and associated with the bottom of the well presumably where the supported lipid bilayers were located (**Fig. 8B, middle panel**).

Here, the VD1 condensates appear to have a slightly increased surface tension (compared to the absence of the SLB), as indicated by the decrease in spreading and the increase in number of distinct surface-wetted areas. In the presence of cardiolipin, there are smaller and fewer spherical condensates floating in solution, while the surface-wetting condensates have increased in number (compared to the absence of cardiolipin), indicating a further increase in surface tension (**Fig. 8B, right panel**). Taken together these data support that VD1 phase separation might present the cardiolipin-binding motif (42) in a coherent manner that would allow for enhanced binding to the membrane.

## Discussion

In this work, the isolated variable domain from Drp1 isoform 1 (VD1) is shown to be intrinsically disordered and remains disordered in the presence of TMAO, a stabilizing osmolyte known to induce folding of proteins with destabilized or latent folds. We show that rather than folding, the variable domain undergoes a cooperative transition to a separate phase, which is also supported by other solutes including the known crowding reagents PEG-8000 and Ficoll PM 70. Phase separation of the variable domain appears to be liquid-like based on FRAP measurements and the ability of droplets to fuse. This liquid-like phase separation (LLPS) enhances membrane binding and is further promoted by the mitochondrial lipid cardiolipin. Reciprocal favorable interaction appears between the membrane and the condensed state suggesting that phase separation may bias the conformational states available to VD1.

By light scattering, both PEG-8000 and Ficoll PM 70 conditions appear to scatter less light than the (NH_4_)_2_SO_4_ condition (**Fig. 6A**), despite a nearly equal number of droplets formed in these conditions (**Fig. 6B-D**). These observations are representative of many replicates under a much wider range of conditions than reported here. The basis for this potential discrepancy is the increased propensity of the PEG-8000 and Ficoll PM 70 conditions to promote the formation of large, amorphous droplets found on the glass bottom of the 96-well plates. This results in larger “base layers” than the (NH_4_)_2_SO_4_ condition and consequently reduced light scattering. This base layer can be seen in the DIC images in Figure 6 and also in the no bilayer control condition in Figure 8B. Pre-coating the plates with passivating agents or surfactants did not prevent base layer droplet formation (**Fig. S6**), nor did the addition of chelators.

A limitation of this study is the requirement for co-solutes for phase separation to occur. However, the co-solutes used here typically do not force proteins into non-native states as they do not preferentially, or directly, interact with the polypeptide backbone or sidechains, and thus do not lower the energy of a particular conformation. Rather, these co-solutes destabilize unfolded conformations to varying degrees by altering the solvation of the protein backbone allowing the protein to move into a lower energy state that is already part of its energy landscape (47, 69–78). The solute-mediated changes in the solvation of VD1 are likely necessary as the N- and C-termini are not constrained in our experiments as they are found in the full-length protein. Despite this, we propose that the phase separation properties are biologically relevant. Mutational analyses of the variable domain dramatically alter Drp1 foci formation and distributions to cause fluid-like wisps of Drp1 by fluorescence microscopy, often non-mitochondrial (see Figure 3 in Reference (31)). These observations are not well explained by current models for Drp1 activity that involve the well-ordered assembly of Drp1 into pre-scission complexes. By contrast, these earlier observations might be explained if Drp1 is able to undergo liquid-liquid phase separation in cells, which is modulated by the VD.

In other phase-separating systems, bridging of like-charges by oppositely charged ions is one mechanism of LLPS. For example, the interaction between arginine and the phosphate backbone of RNA is observed in many phase-separating systems (79), and polyArg has been shown to phase separate with a variety of polyanions, including (but not limited to) RNA, DNA, chondroitin, heparin, and polyphosphate

(80). Cardiolipin has a unique diphosphate headgroup, and therefore a cardiolipin-rich membrane is essentially a two-dimensional polyphosphate. The observation of VD phase separation in the presence of TMAO and ammonium sulfate may be due in part to TMAO and sulfate anion acting as a proxy of sorts for cardiolipin. In other words, cardiolipin-Arg interactions may screen repulsive Arg-Arg forces to allow intermolecular VD-VD interactions at the membrane that enhance phase separation. Consistent with this idea, an Arg-rich motif of RRNRLAR exists in VD1 and prior work mutating the RRNR to AANA prevented Drp1 from remodeling cardiolipin-containing vesicles and reduced Drp1 puncta formation in cells. Consistent with the VD ability to form different phases, previous work has shown that nanometer-scale clustering of cardiolipin is induced by transient VD-membrane interactions that can subsequently coalesce into condensed regions (11). A prediction from these considerations is that the AANA substitution in VD1 would reduce its propensity to form LLPS in the presence of cardiolipin-containing membranes.

A recently identified cardiolipin-binding motif in VD (Trp-Arg-Gly) when deleted impaired fission indicating an important role for VD-cardiolipin interactions in fission (42). In our MD simulations of VD1, this tryptophan is solvent-exposed (**Fig. 3C-D**) that likely is necessary for engaging cardiolipin. We suspect that the phase-separated states bias this conformation over others to enhance lipid binding. VD1-modulated phase separation of Drp1 could allow for rapid, tunable assembly of the polymer necessary for fission. In this model, liquid droplets composed of Drp1 (and possibly other interactors) provide a high local concentration of protein to overcome the nucleation barrier for polymerization, which could be tuned by the numerous post-translational modifications (or alternative splice variants) that occur in the VD to govern Drp1 activity. Indeed, such droplet-mediated assembly of microtubules also occurs, where tau initially forms droplets in the presence of crowders, then recruits tubulin, concentrating it within the droplet and enabling polymerization of the tubulin subunits into stable microtubules (81).

## Experimental Procedures

### Drp1 and Drp1ΔVD1 cloning

Drp1 isoform 1 (Genbank accession number AB006965) was PCR amplified with Pfu Turbo DNA polymerase (Stratagene, La Jolla, CA) as an NdeI/XhoI fragment, with a Tobacco Etch Virus (TEV) protease site (ENLYFQS) preceding the XhoI restriction site. The resulting DNA fragment was then ligated into the bacterial expression vector pET29b (EMD Millipore, Billerica, MA), which contains a C-terminal hexa-histidine (His6) tag. To clone a Drp1ΔVD-His6 fusion protein, the gene sequence corresponding to residues 1-525 of Drp1 isoform 1 was fused to the gene sequence corresponding to residues 637-736 of Drp1 isoform 1 with a “GGGSGGG” linker. The Drp1ΔVD-His6 fusion protein was created by PCR amplifying two fragments. The first fragment contained an NdeI restriction site, the Drp1 GTPase domain (residues 1-525), and the “GGGSGGG” linker. The second fragment contained the same linker, residues 637-736 of the GED domain, a TEV cleavage site (ENLYFQS) and an XhoI restriction site. 100 ng of each fragment were mixed together to form the template of a third PCR reaction, which was amplified with the forward primer of reaction 1 and the reverse primer of reaction 2. The resulting 1100 bp PCR product was gel extracted, digested with NdeI/XhoI and ligated into pET29b.

### Drp1 and Drp1ΔVD1 expression and purification

pET29b-Drp1-His6 or pET29b-Drp1ΔVD-His6 were transformed into Escherichia coli BL21 (DE3). Cells were grown at 37°C in Super Broth (SB) with kanamycin (30 μg/ml) to an A600 of ∼1.5 with shaking at 250 rpm, the temperature was lowered to 14°C and after 30 minutes, protein expression was induced with 0.5 mM isopropyl 1-thio-β-D-galactopyranoside for 12-16 hours. Cells were harvested by centrifugation using a Sorvall JLA-8.1000 rotor at 5,000 rpm for 10 minutes at 4°C and were resuspended in column buffer A (50 mM 4-(2-hydroxyethyl)-1-piperazineethanesulfonic acid (HEPES), pH 7.4, 400 mM NaCl, 5 mM MgCl_2_, 40 mM imidazole) containing protease inhibitors (Complete, EDTA-free Protease Inhibitor Mixture, Roche Applied Science, Indianapolis, IN). Cells were lysed by four passes through a French press (Thermo Scientific, Pittsburgh, PA), DNAse was added to a final concentration of 1 μg/ml and lysates were clarified by centrifugation using a Sorvall SS34 rotor at 15,000 rpm for 30 minutes at 4°C. His6-tagged fusion proteins were isolated from the resulting supernatant by affinity chromatography using Ni^2+^ Sepharose High-Performance beads (GE Healthcare, Pittsburgh, PA). Bound fusion proteins were washed with 200-300 mL of column Buffer A, 100 mL of column buffer B (50 mM HEPES, pH 7.4, 400 mM NaCl, 5 mM MgCl_2_, 40 mM imidazole, 1 mM ATP, 10 mM KCl), 200 mL of column buffer C (50 mM HEPES, pH 7.4, 80 mM imidazole, 400 mM NaCl, 0.5% (w/v) 3-[(3-cholamicopropyl)dimethylammonio]-1-propanesulfonate (CHAPS)), and 100 mL of column buffer A. Bound fusion proteins were then eluted with column buffer D (50 mM HEPES, pH 7.4, 800 mM NaCl, 5 mM MgCl_2_, 500 mM imidazole). Fractions containing His6-tagged fusion proteins were pooled and dialyzed overnight at 4°C into buffer containing 50 mM HEPES, pH 7.4, 5 mM MgCl_2_, 1 M NaCl with Spectra/Por Biotech Cellulose Ester dialysis membrane with a 100,000 molecular weight cut-off (Spectrum Laboratories, Rancho Dominguez, CA). Proteins were concentrated by centrifugation in Vivaspin 20 ultrafiltration devices with a 100,000 molecular weight cut-off (GE Healthcare). Protein concentrations were quantified by UV spectroscopy with an extinction coefficient of 36380 M^-1^cm^-1^ (Drp1) or 30870 M^-1^cm^-1^ (Drp1ΔVD) after incubation for 3 hours in 6M guanidinium hydrochloride at 65°C. Purified protein was stored at 4°C until use and was used with 72 hours of cell lysis.

### VD1 cloning, expression and purification

Boundaries of the VD1 were identified by performing a multiple sequence alignment with classical dynamins and dynamin-related proteins and identifying the interruption of sequence conservation. Additionally, the Drp1 amino acid sequence was analyzed using the JPRED structure prediction algorithm to identify where the regular secondary structure of the stalk domain was predicted to be interrupted. This prediction was consistent with the multiple sequence alignment. Finally, a homology model was constructed using Swiss-Model, with the dynamin-1 crystal structure as a template, and the interruption of the helical stalk domain was again identified. A high-resolution crystal structure for Drp1 (with the VD1 removed) has since been solved, and the termination of helices in the stalk domain at either terminus of the VD1 observed in the crystal structure agrees with our multiple sequence alignment, JPRED structure prediction and homology model. A VD1 construct representing amino acids 501 – 637 of human Drp1 isoform 1 was PCR amplified from a full-length Drp1 template as Nde1/Xho1 fragment with a Tobacco Etch Virus (TEV) protease site (ENLYFQS) preceding the XhoI restriction site, and subcloned into the pET29b expression vector (EMD Biosciences) including a C-terminal 6xHis tag. All constructs were verified by DNA sequencing (GENEWIZ, South Plainfield, NJ). Plasmids were transformed into chemically competent Escherichia coli Rosetta cells (Novagen) and grown at 37°C in Super Broth with kanamycin (30mg/mL) and chloramphenicol (34mg/mL) to A600 of 1.0. Protein expression was induced by addition of 0.5 mM isopropyl ß-D-1-thiogalactopyranoside (IPTG) at 18°C and harvested by centrifugation 15–18 h after induction. The resulting cell pellets were resuspended in Ni column buffer (25 mM Tris HCl, 50 mM NaCl, 30 mM imidazole pH 7.4) containing protease inhibitors (Roche Applied Science). Cells were lysed with 4 passes through an Emulsiflex C3 (Avestin), DNase was added to 1 mg/mL and lysates were clarified by centrifugation using a Sorvall SS34 rotor at 15,000 rpm for 30 minutes at 4°C. Protein was isolated from the resulting supernatant by affinity chromatography using Ni-Sepharose 6 Fast Flow beads (GE Healthcare), and eluted with a 100 mL linear gradient of column buffer with 500 mM imidazole. Fractions containing His-tagged proteins were pooled, TEV protease was added at 1:100 molar ratio, and the solution was dialyzed against SP column buffer (50 mM KPhos, 50 mM KCl, 1 mM DTT, pH 7.4) using Spectra/Por Biotech Cellulose Ester dialysis membrane with a 12 – 14 kDa molecular weight cut-off (Spectrum Laboratories, Rancho Dominguez, CA) at 4°C for 24 hours or until the protease reaction reached completion, as determined by SDS-PAGE. VD1 was separated from TEV protease and further purified on a HiTrap SP XL column (GE Healthcare). Flow-through fractions containing VD1 were pooled and concentrated to ∼ 1 mM by centrifugation in Amicon Ultra-15 ultrafiltration devices with a 10 kDa molecular weight cut-off (EMD Millipore, Billerica, MA). Sample purity was checked by Coomassie-stained SDS-PAGE and was typically greater than 95%. Protein concentrations were determined by UV spectroscopy using a molar extinction coefficient of 6990 M^−1^cm^−1^. Concentrated protein stocks were divided into 50 μL aliquots, flash frozen in liquid nitrogen, lyophilized and stored under dry conditions at −20°C until use. Immediately prior to use, lyophilized stocks were reconstituted using 50 μL of deionized water, and protein concentration was re-measured.

### Size Exclusion Chromatography (SEC)

VD1 hydrodynamic properties were approximated by SEC using a HiLoad 16/60 Superdex 75 prep grade (S-75) column (GE Healthcare) in 20 mM Tris, 50 mM NaCl, 1 mM DTT, pH 7.4.

### Sedimentation velocity AUC

Sedimentation velocity experiments were carried out in a Beckman XL-A analytical ultracentrifuge, using two-sector cells and an An60Ti rotor. Experiments were carried out at a speed of 50,000 rpm and 22°C. Sedimentation profiles were detected using absorbance optics operated in continuous mode. The sedimentation coefficient, apparent diffusion coefficient and MW were determined by fitting data to the Lamm equation using the DCDT+ software (40, 82, 83).

### NMR spectroscopy

NMR experiments were performed at 18.8 T on a Varian Inova 800 spectrometer outfitted with a TXI coldprobe. Two-dimensional ^1^H–^15^N heteronuclear single quantum correlation (HSQC) experiments were collected at pH 7.4 using WATERGATE for solvent suppression. Uniformly ^15^N-labeled VD1 samples were prepared by dialysis into 20 mM Tris-HCl pH 7.4. with 100 mM NaCl, 10 mM EDTA, 1 mM DTT and 10% D2O. The protein concentration was 300 μM. HSQC spectra were collected at 4 scans per increment, 1280 (t2) x 256 (t1) complex points with acquisition times of 64 ms (^1^H) and 71 ms (^15^N). Carrier frequencies were centered on the chemical shift of water in ^1^H and in the center of the amide region at 117.5 ppm in ^15^N. NMR data were processed using NMRPipe (84) and analyzed with NMRView (85). For spin relaxation experiments, R_2_ spin relaxation experiments were collected at 298K on a sample of ^15^N-VD1 using a standard CPMG pulse sequence at 18.8 T. The T_2_ delays of 17.6, 35.2, 52.8, 88.0, 123.2, 158.4 ms were collected in random order to reduce systematic errors with two time points recorded in duplicate for error analysis. Peak heights were extracted and analyzed using the NMR Series tool in CCPN NMR Analysis software (86) (https://ccpn.ac.uk/software/version-2/). Spin relaxation rates were determined by nonlinear least-squares optimization tool in NMR Analysis to fit data to a single exponential for each residue.

### Preparation of TMAO buffers

Trimethylamine N-oxide dihydrate (98% pure; Sigma-Aldrich) was dissolved in 100 mM Tris, 200 mM NaCl, 50 mM arginine buffer to make 0, 0.8, 1.6, 2.4, 3.2, 4.0 and 4.8 M TMAO buffers. The pH was adjusted to 7.4 for each buffer separately. Impurities in the TMAO were removed by incubating each buffer solution with activated carbon (12–20 mesh; Sigma-Aldrich) for at least 4 hours while protected from light. The buffer was then filtered (0.22 μm filter; Millipore), aliquoted, flash frozen in liquid nitrogen and stored at −80 °C until further use. At the time of sample preparation, TMAO buffers with similar concentrations were mixed in the appropriate ratios to obtain the target TMAO concentrations used in the fluorescence measurements, which was necessary to minimize pH changes that can occur when mixing two TMAO buffers with a large concentration difference. Final TMAO concentrations were determined by refractive index and a standard curve, after the manner of Bolen (87).

### Circular dichroism

Far UV circular dichroism spectra for the Drp1 isoform 1 VD1 were recorded with an Aviv Model 215 CD spectrometer (Aviv Biomedical, Lakewood, NJ) from 260 nm to 195 nm with a bandwidth of 1.0 nm and scan step of 1 nm in a 0.1 cm quartz cuvette at 22°C. All spectra were recorded in CD buffer (10 mM sodium phosphate, 200 mM sodium fluoride, 1 mM TCEP, pH 7.4) and corrected for the contribution of buffer. Each spectrum shown is an average of three spectra. Data for which the HT voltage rose above 500 V were discarded. Mean residue ellipticity ([θ], deg cm^2^/dmol/res) was calculated using the equation [θ] = θ × *MRW*/(10×0. 1 *cm*×[*B*]), where *MRW* = *MW*/(# *residues* − 1). CD spectra were deconvoluted using CONTIN-LL in the CDPro software package (11, 32).

### Steady-state tryptophan fluorescence

Steady-state fluorescence emission spectra of the VD1 were measured using an Aviv ATF-105 fluorometer (Aviv Biomedical, Lakewood, NJ) in TMAO buffers of varying concentrations. Fresh dithiothreitol (DTT) was added to a final concentration of 15 mM to a 15x protein stock solution, to be diluted to 1mM DTT upon sample preparation. Protein samples (ranging from 2 μM to 40 μM) were prepared in a final volume of 150 μL by combining 10 μL of 15x protein stock with 140 μL of TMAO buffer of the appropriate concentration. Samples were allowed to equilibrate in a “submicro” fluorimeter cell at 22 °C (Starna, Atascadero, CA) for 5 min to allow for temperature stabilization and protein conformation equilibrium to be reached. Emission spectra were then recorded with excitation at 295 nm (5 nm slit width). All spectra were corrected for the contribution of buffer. Fluorescence emission intensities at 338 nm (5 nm slit width) were recorded as a function of TMAO concentration. The resulting sigmoid curve was fit to a two-state cooperative folding transition using nonlinear least-squares analysis to determine the stability (ΔG) and m-value (88, 89). In order to maximize the amount of spectral information used for analysis, we calculated the center of spectral mass 〈ν〉 from fluorescence spectra using equation 1:

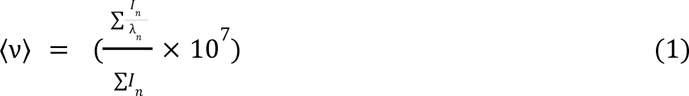

where λ is the wavelength in nm, *I_n_* is the fluorescence emitted at wavelength λ*_n_*, and the summation is carried out from λ*_n_* = 310 *nnm* to λ*_n_* = 480 *nnm*. The variable *v* denotes the wavenumber, defined as 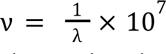, and has been used historically because it is directly proportional to energy. It has been shown that the center of spectral mass does not scale proportionally with fraction folded protein (90), but this can be corrected for when determining thermodynamic values from these data (91).

### Light scattering

Right-angle light scattering near 350 nm was measured using an Aviv ATF-105 fluorometer (Aviv Biomedical, Lakewood, NJ) immediately following each fluorescence measurement. Excitation and emission wavelengths were set to 345 and 355 respectively, with both slit widths at 5 nm. Excitation and emission wavelengths had to be offset in this manner in order to keep the signal in a measurable range without altering the PMT settings used for fluorescence measurements. Additionally, samples were retained after fluorescence measurements and their absorbance spectra were measured with a Nanodrop 2000c UV/visible spectrophotometer (Thermo Scientific) in a quartz cuvette with a 1 cm pathlength. The absorbance at 350 nm was used as an indicator of scattered light since the VD1 does not absorb light at this wavelength under native conditions. The two methods were found to give comparable results. Light scattering data were fit to a two-state model after the same manner as outlined above for fluorescence data.

### Electron microscopy

Samples were prepared for EM using samples retained from fluorescence measurements, or with fresh samples prepared in the same manner. 5 μL of protein sample was applied to freshly-ionized carbon-coated copper mesh grids (Electron Microscopy Science). After 15 minutes, the grids were rinsed quickly with four washes of deionized water, then floated face-down on a drop of 2% uranyl acetate solution for one minute before being air-dried. Images were taken on a FEI Tecnai 12 TWIN equipped with 16 bit 2K x 2K FEI Eagle bottom mount camera and SIS Megaview III wide-angle camera (Olympus).

### Vesicle Extrusion

All synthetic lipids were obtained from Avanti Polar Lipids (Alabaster, AL). Lipids were measured from chloroform stocks using Hamilton syringes, mixed in the intended ratios, and dried in a thin film under a nitrogen stream. Excess chloroform was removed from dried films by lyophilization for at least 2 hours. The dried lipids were then resuspended in the appropriate amount of deionized water to make a 13.1 mM solution of lipids. Lipid solutions were subjected to 11 freeze-thaw cycles in a dry ice/ethanol bath and a 37°C water bath. Freeze-thawed solutions were extruded at least 31 times through a 100 nm nucleopore track etch membrane (Whatman), using an Avanti syringe extruder apparatus (Avanti Polar Lipids). Vesicle size, homogeneity and reproducibility were verified using dynamic light scattering and electron microscopy. All lipids had dioleoyl (DO) acyl chains with the exception of cardiolipin, which had tetraoleoyl (TO) acyl chains. Vesicles used for sedimentation included 0.25% 1,2-dioleoyl-sn-glycero-3-phosphoethanolamine-N-(lissamine rhodamine B sulfonyl) (Rh-DOPE) for easy visualization of lipid pellets.

### Calculation of partition coefficients and free energy of partitioning

The lipid partition coefficient was calculated using equation 3 (92, 93) from binding data measured as a function of LUV composition (increasing fraction of acidic lipid in a DOPC background):

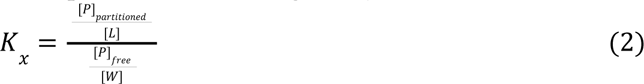

The concentration of protein partitioned, [*P*]_partitioned_ = *f_p_*x[*P*]_*total*_, and the concentration of free protein, [*P*]_*free*_= [*P*]*_total_*− [*P*]_*partitioned*_. [*L*] is the accessible acidic lipid concentration, taken to be half of the total acidic lipid concentration to account for the inaccessible lipids on the inner leaflet of the lipid bilayer. The concentration of water, [*W*] was considered to be 55.3 M.

Using the partition coefficients obtained from equation 2, a free energy of lipid partitioning was obtained from equation 4:

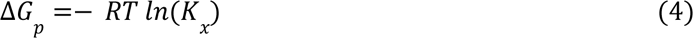

*R* is the gas constant in kcal/mol T, and *T* is temperature in Kelvin. Since only half of the total acidic lipid concentration was used for calculations of *K_x_* (described above), a correction factor of −0.41 kcal/mol was added to the calculated Δ*G_p_* as suggested (92).

### MD simulations

Initial VD1 structures were constructed in random coil configurations, placed in a solvent box of size 72Å x 72Å x 72Å with a water-ion solution containing 0.2 M NaCl, and equilibrated for 300 ns. Similarly, single solvated VD1 with TMAO was constructed and run for 300 ns to investigate the effect of TMAO. All simulations were carried out using NAMD 2.10 (94, 95) with a time step of 2 fs and a Langevin thermostat and barostat were employed under NPT conditions at 300 K and 1 atm. The van der Waals (vdW) interaction cut-off was set to 1.2 nm and electrostatic interactions were computed using the particle-mesh Ewald (PME) algorithm. The CHARMM36 force field (96) with the protein backbone improvement (97) and dihedral angles optimization (98) was adopted for the protein and the TIP3P model was used for the water molecules.

### Fluorophore labeling of VD1

Purified VD1 was dialyzed into SP Buffer (50 mM KPhos pH 7.4, 50 mM KCl, 1 mM TCEP-HCl). AlexaFluor 568 C_5_ maleimide (Thermo) was added at a of 1:20 molar ratio of VD1:fluorophore. Conjugation reactions were carried out by gently rocking for over 12 hours at 4°C protected from light. Free unconjugated dye was removed from labeled VD1 with a Sephadex G-25 M PD-10 desalting column (GE) eluted with pure water. Labeled protein was aliquoted and lyophilized, then later reconstituted in pure water and stored at −20°C. Protein concentration and degree of labeling were determined via Nanodrop 2000c UV/visible spectrophotometer (Thermo).

### Phase separation assays

Purified VD1 was dialyzed into HEPES buffer (20 mM HEPES pH 7.4, 1 mM beta-mercaptoethanol), then aliquoted and stored at −20°C until use. VD1 (1% AlexaFluor 568 C_5_ maleimide-labeled) was added to wells of a glass bottom 96-well plate (Cellvis) containing HEPES pH 7.4, KCl, and pure water. This mixture was briefly mixed and incubated for 5 minutes at room temperature protected from light. Phase separation was induced upon the addition of either (NH_4_)_2_SO_4_, PEG 8000 (Sigma), or Ficoll PM 70 (Sigma) to the reaction mixture and thoroughly mixed. Total reaction volume was 100 μL across all experiments. Reagent volumes and concentrations varied depending on desired conditions. 96-well plates were sealed tightly with parafilm and stored at room temperature protected from light for at least 24 hours prior to downstream application.

### Phase separation light scattering

Absorbance at 340 and 540 nm was measured using a FlexStation 3 Multi-Mode Microplate Reader (Molecular Devices) immediately following each fluorescence measurement. The absorbance at both 340 and 540 nm were used as indicators of scattered light since VD1 does not absorb light at these wavelengths under native conditions. Light scattering data were baseline corrected with buffer-matched controls and plotted using the R Statistical Package (R Core Team, 2013).

### Fluorescence microscopy

Differential interference contrast (DIC) and confocal fluorescence microscopy images were captured on a Nikon Eclipse Ti microscope base equipped with a Yokogawa CSU-W1 spinning disk confocal scanner unit (50 µm pinhole), 40x 0.90 NA objective, and Hamamatsu ORCA-Flash4.0 V3 sCMOS camera. Representative regions of each well were captured with 2x averaging enabled. DIC images were captured after a 1 second exposure. Fluorophore activation was stimulated using a 561 nm laser at 50% power, and images were captured after a 0.200 second exposure.

### FRAP microscopy

Fluorescence recovery after photobleaching (FRAP) was achieved using a Nikon A1R laser scanning confocal microscope equipped with a Galvano resonant scanner unit and 100x 1.45 NA objective. A 54.6028 by 3.4127 µm rectangle was imaged with the 100x objective (zoom 2.331; 512x32 pixels scanning range). A 0.4 µm diameter circle placed within a droplet was photobleached for 0.114 s with a laser power of 15%, 30%, or 100% for droplets in (NH_4_)_2_SO_4_, PEG 8000, or Ficoll PM 70, respectively. Twenty images were taken prior to photobleaching, and two hundred forty-four images were taken after photobleaching. For droplets in (NH_4_)_2_SO_4_, images were captured every 0.160 seconds for a combined elapsed time of 42.24 seconds. For droplets in PEG 8000 and Ficoll PM 70, images were captured every 0.140 seconds for a combined elapsed time of 36.96 seconds. Using the Nikon Advanced Research software, three regions were measured: FRAP stimulation, nonspecific photobleach, and background. The data were normalized, background subtracted, and bleach corrected using the FRAP analysis tool in the software package. Conditions were setup in an 18-chambered #1.5 coverglass system (C18-1.5H; Cellvis) plate and droplets in (NH_4_)_2_SO_4_, PEG 8000, and Ficoll PM 70 were imaged 24, 44, and 44 hours after assay setup.

### Image Processing

Image analysis was performed using ImageJ (FIJI; Versions 1.52n and newer). Equivalent LUTs ranges were used for depicting AlexaFluor 568 fluorescence in all conditions across all experiments. Roughly equivalent brightness and contrast settings were used for DIC images (slightly adjusted for clarity and visual acuity). Microscopy data processed by ImageJ was plotted using the R Statistical Package.

### Phase diagram

Varying concentrations of potassium chloride were titrated against a concentration gradient of VD1 (1% AlexaFluor 568 C_5_ maleimide-labeled) following phase separation assay setup as outlined previously and imaged using DIC and confocal fluorescence microscopy. Each well was acquired with identical microscopy settings. Representative regions and focal planes of each image were selected, for which mean pixel intensity and standard deviation were calculated in ImageJ. Resultant .tif files were baseline corrected with the standard deviation of intensity from a no fluorophore control condition. Coefficient of variation values for each condition were determined then plotted using the R Statistical Package.

### Supported lipid bilayer (SLB) preparation

All synthetic lipids were obtained from Avanti Polar Lipids. MitoMix (+CL) lipid mixture: 48% egg PC (PC; L-α-phosphatidylcholine; 840051), 28% 16:0-18:1 PE (POPE; 1-palmitoyl-2-oleoyl-sn-glycero-3-phosphoethanolamine; 850757), 10% soy PI (PI; L-α-phosphatidylinositol; 840044), 10% 18:1 PS (DOPS; 1,2-dioleoyl-sn-glycero-3-phospho-L-serine; 840035), 4% cardiolipin (CL; 1’,3’-bis[1,2-dioleoyl-sn-glycero-3-phospho]-glycerol; 710355). MitoMix (-CL) lipid mixture: 44% PC, 28% POPE, 10% PI, 18% DOPS. To prepare large unilamellar vesicles (LUVs), MitoMix and MitoMix (-CL) lipid mixtures were dissolved in chloroform. The solvent was allowed to evaporate for over 12 hours in open air, at which point residual chloroform was dried under nitrogen gas. LUVs were resuspended in an appropriate volume of buffer (20 mM HEPES pH 7.4, 500 mM NaCl, 40 mM Imidazole, 0.02% NaN_3_) to achieve a 10 mM solution. Lipid solutions were subjected to 11 freeze-thaw cycles in a dry ice/ethanol bath and a 37°C water bath. Freeze-thawed solutions were extruded at least 21 times through a 100 nm nucleopore track etch membrane (Whatman), using an Avanti syringe extruder apparatus (Avanti Polar Lipids). A glass bottom 96-well plate (Cellvis) was treated with ozone for 1 hour, at which point a 1mM solution of appropriate lipid mixture was added such that the bottoms of the desired wells were covered. The plate was incubated at a temperature higher than the phase transition temperature of the lipids for 1 hour to form an SLB, then allowed to cool at room temperature for 30 minutes. Wells were thoroughly rinsed with phase separation buffer (10 mM HEPES pH 7.4, 10 mM KCl) to remove residual lipids and immediately prepared for phase separation assays. Final buffer conditions for these experiments were 10 mM HEPES pH 7.4, 50 mM KCl, and 20% PEG 8000.

All custom code is available from the authors upon request.

## Supporting information

Posey et al supplemental information

## Acknowledgments

We thank Dr. John A Corbett for the generous use of his laboratory’s confocal microscope, Dr. Rajesh Ramachandran for helpful discussions and sharing unpublished data, and the Cardiovascular Center at MCW for allowing use of their Nikon A1R microscope for FRAP imaging. This work was supported by the National Institutes of Health grants: R01-GM63747 (VJH) and R01-GM067180 (RBH), and by a Discovery grant RGPIN-2018-06886 from the the Natural Sciences and Engineering Research Council of Canada (JLH).

## CRediT author statement

**AEP** - Conceptualization, Data curation, Formal Analysis, Investigation, Methodology, Validation, Visualization, Writing

**MB** - Conceptualization, Data curation, Formal Analysis, Investigation, Methodology, Validation, Visualization, Writing

**KAR** - Conceptualization, Data curation, Formal Analysis, Investigation, Methodology, Validation, Visualization, Writing

**ENL -** Data curation, Formal Analysis, Investigation, Validation, Writing – review & editing

**MAK** - Data curation, Formal Analysis, Investigation, Validation, Writing – review & editing

**CMJ** - Data curation, Investigation, Methodology, Validation, Writing – review & editing

**MCH** - Conceptualization, Data curation, Formal Analysis, Investigation, Methodology, Supervision, Validation, Visualization, Writing – review & editing

**NWK** - Data curation, Formal Analysis, Investigation, Validation, Visualization, Writing – review & editing

**VJH** - Conceptualization, Project administration, Supervision, Validation, Visualization, Writing – review & editing

**JLH** - Conceptualization, Data curation, Formal Analysis, Funding acquisition, Investigation, Methodology, Project administration, Resources, Supervision, Validation, Visualization, Writing

**RBH** - Conceptualization, Data curation, Formal Analysis, Funding acquisition, Investigation, Methodology, Project administration, Resources, Supervision, Validation, Visualization, Writing

