## Supplementary material for "The variable domain from the mitochondrial fission mechanoenzyme Drp1 promotes liquid-liquid phase separation": Posey et al supplemental information

**Supplemental Information**

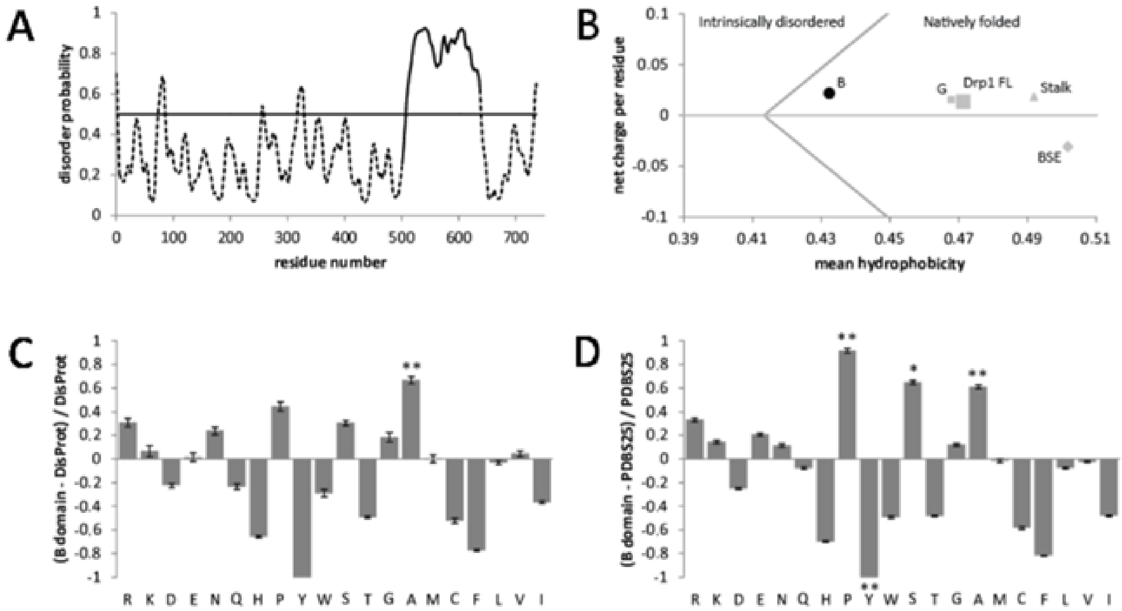


E,F

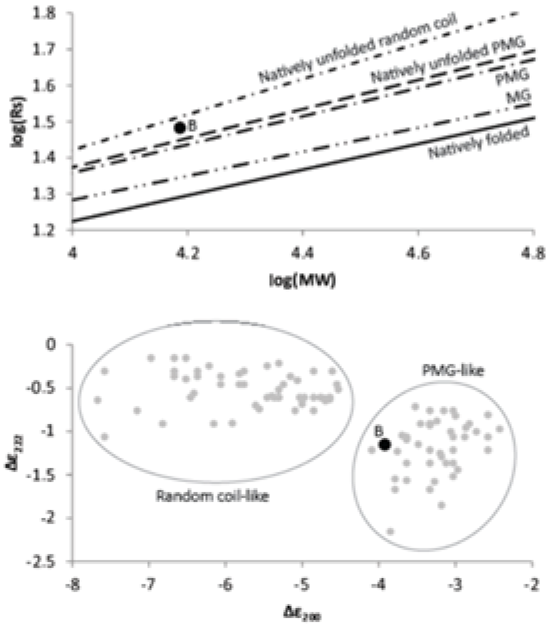


**Supplemental Figure 1. Properties of the Drp1 variable domain compared to other proteins with intrinsic disorder.** (A) PrDOS disorder probability prediction for Drp1. (B) Net charge vs. hydrophobicity plot for full-length Drp1 (Drp1 FL) and the individual Drp1 domains. (C,D) Amino acid composition profiles of variable domain vs. the DisProt database or a selection of globular proteins from the PDB. (E) The Stokes radius (Rs) of the variable domain (filled black circle) lies between that of a natively unfolded pre-molten globule (PMG) and a natively unfolded coil, according to standard curves published by Uversky (39, 41). (F) The molar ellipticities from circular dichroism measurements of the variable domain at 200 and 222 nm are similar to PMG-like standards published by Uversky (39, 41). ** significance level α = .05, * significance level α = .051. All measurements at 25°C.

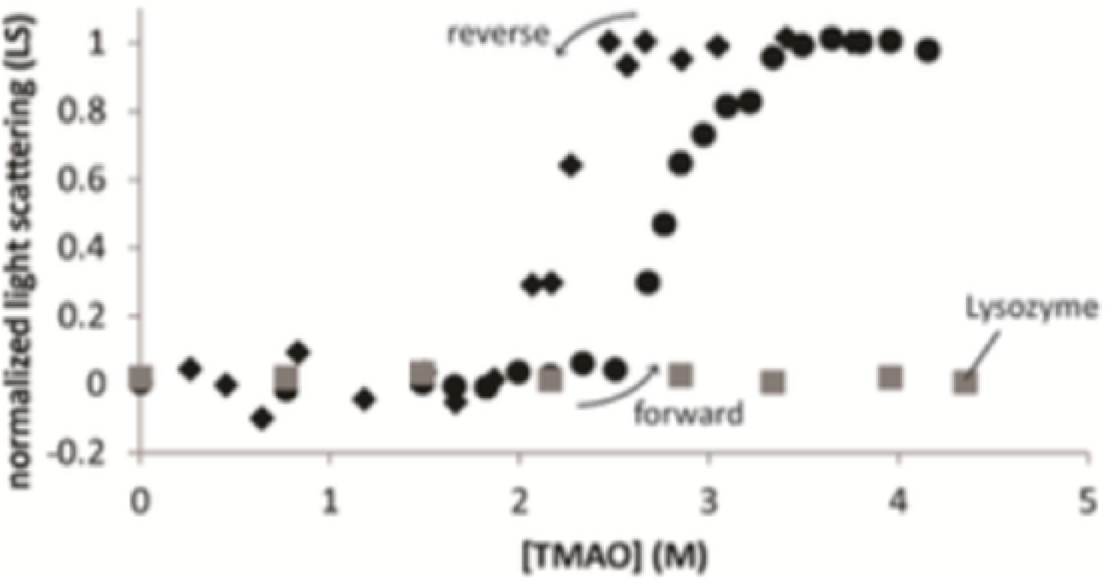


**Supplemental Figure 2. TMAO titrations with lysozyme.** TMAO titration of the variable domain followed by light scattering gives a two-state transition similar to that observed by fluorescence, with similar hysteresis between forward (filled circles) and reverse (filled diamonds) titrations. Lysozyme, used here as a negative control, does not scatter light in TMAO (gray squares).

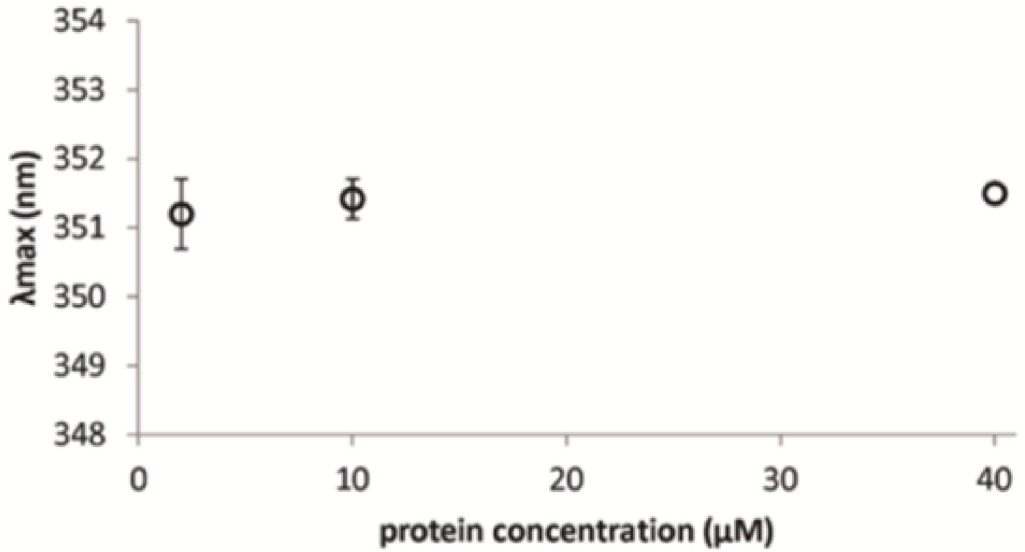


**Supplemental Figure 3. Concentration dependence of the variable domain intrinsic tryptophan fluorescence.** Emission spectra were recorded and $\lambda_{max}$ is plotted at indicated concentrations of isolated variable domain.

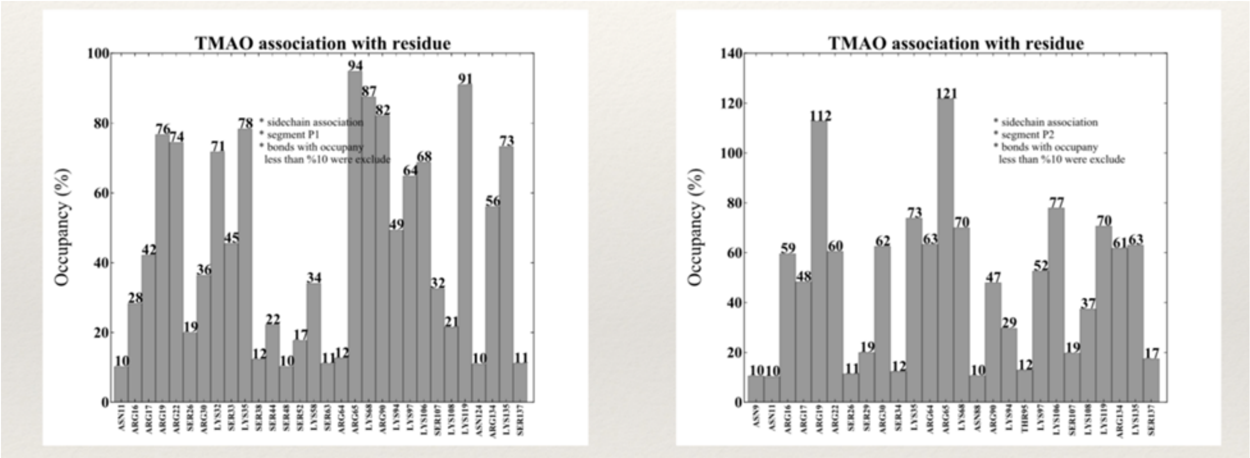


**Supplemental Figure 4. TMAO interactions with the variable domain.** To investigate dominant TMAO interactions, protein residues that were in contact with TMAO for more than 10% of the simulation time were recorded, along with their total occupancy with TMAO molecules.

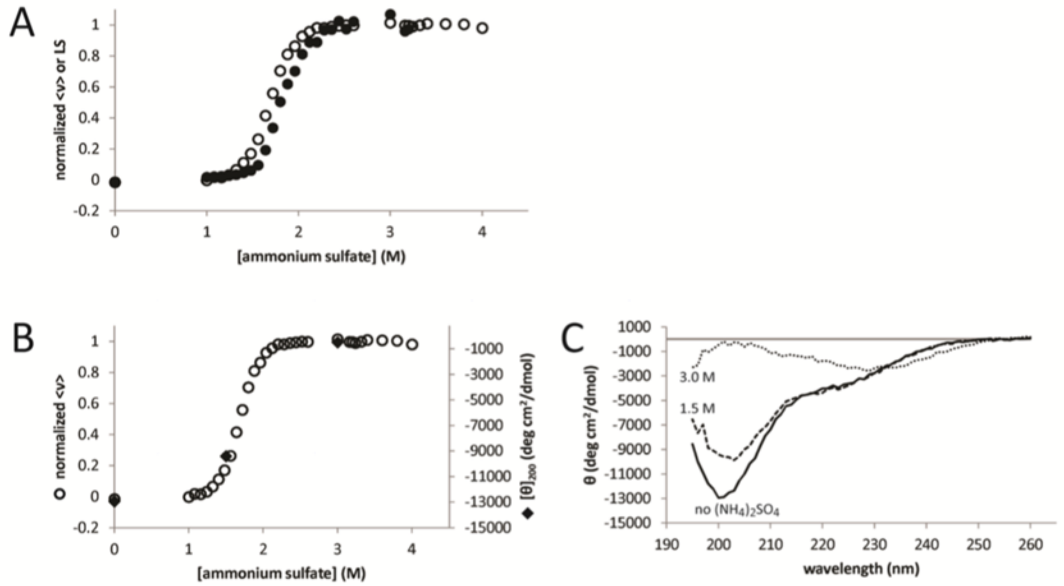


**Supplemental Figure 5. Steady-state tryptophan fluorescence, light scattering and circular dichroism of VD1 in the kosmotropic salt, ammonium sulfate.** (A) The variable domain exhibits a two-state transition by fluorescence (open circles) and light scattering (LS) (filled circles), similar to what was observed in TMAO, except that the light-scattering is slightly shifted to higher ammonium sulfate concentrations. (B) Molar ellipticity of the variable domain at 200 nm (filled diamonds) aligns with the transition observed by fluorescence (open circles). Both measurements were made at 2 μM protein concentration. (C) CD spectra of the variable domain in increasing concentrations of ammonium sulfate ((NH4)2SO4).

To alleviate the problem of optical interference by TMAO, and also to determine whether the VD1 would exhibit similar behavior in the presence of other cosolvents, we repeated fluorescence, RALS and CD measurements of the VD1 in the kosmotropic salt, ammonium sulfate. The VD1 exhibited similar two-state-like behavior as measured by fluorescence and light scattering, although the midpoint of the light scattering curve was slightly shifted toward higher ammonium sulfate concentration compared to the fluorescence curve. This may reflect a slight kinetic lag in assembly/aggregation of the VD1 in ammonium sulfate. The mean residual ellipticity at 200 nm in increasing concentrations of ammonium sulfate overlays with the fluorescence curve (Figure S5A), indicating that these observables are reporting on the same process or concomitant processes. The full CD spectra in ammonium sulfate follow a similar trend as in TMAO, with a minimum appearing near 230 nm, but we were also able to observe a maximum around 200 nm since the absence of optical interference in ammonium sulfate buffers allowed us to make measurements at much lower wavelengths (Figure S5C).

The VD1 also exhibited a response similar to what was observed in TMAO and ammonium sulfate in another protecting osmolyte, sarcosine. As a function of increasing sarcosine concentration, we observed an apparent two-state-like change in mean residual ellipticity that was superimposable on the normalized fluorescence curve (data not shown). Taken together, these data suggest that the observed folding and self-assembly behavior is intrinsic to the VD1 and is independent of kosmotropic cosolvent. As the

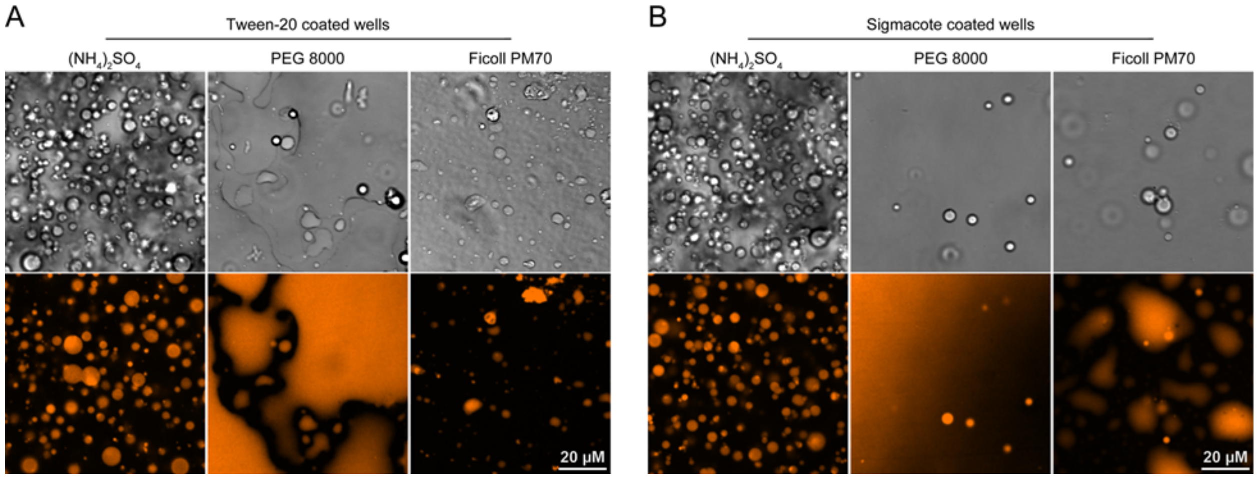


quality of the solvent decreases (due to increasing kosmotropic cosolvent), the VD1 undergoes changes in secondary structure with concomitant assembly/aggregation.

**Supplemental Figure 6. Treatment with surfactants or passivating agents does not inhibit Drp1 VD1 "base layer" formation under LLPS conditions.** DIC and fluorescence microscopy images of 50 µM VD1 in 1.68 M $(NH_4)_2SO_4$ (left), 20% PEG 8000 (middle), and 20% Ficoll PM 70 (right) on either Tween-20-coated (A) or Sigmacote-coated (B) wells.
